# Absence of pathogenic mutations in CD59 in chronic inflammatory demyelinating polyradiculoneuropathy

**DOI:** 10.1101/459917

**Authors:** Lena Duchateau, Lorena Martin-Aguilar, Cinta Lleixà, Andrea Cortese, Oriol Dols-Icardo, Laura Cervera-Carles, Elba Pascual-Goñi, Jordi Diaz-Manera, Ilaria Calegari, Diego Franciotta, Ricard Rojas-Garcia, Isabel Illa, Jordi Clarimon, Luis Querol

**Author notes:** These authors contributed equally (Lena Duchateau, Lorena Martin-Aguilar). Corresponding author: Luis Querol, Hospital de la Santa Creu I Sant Pau, Universitat Autònoma de Barcelona, Mas Casanovas, 90, 08041, Barcelona, Spain.

## Abstract

**Objective:** Mutations in *CD59* cause CIDP-like polyneuropathy in children with inherited chronic hemolysis. We hypothesized that mutations in *CD59* might be found in a subset of sporadic CIDP patients.

**Methods:** 5 patients from two centers, fulfilling the EFNS/PNS diagnostic criteria for CIDP were included. *CD59* coding region was amplified by PCR and Sanger sequenced.

**Results:** One rare variant was detected in a patient which resulted in a synonymous change and predicted to be neutral. Pathogenic variants were absent in our cohort.

**Interpretation:** Our pilot study suggests that mutations in *CD59* are absent in adult-onset sporadic CIDP.

## INTRODUCTION

Chronic inflammatory demyelinating polyradiculoneuropathy (CIDP) is a rare and heterogeneous neurological disorder that is diagnosed according to clinical and electrophysiological diagnostic criteria. Its pathogenesis is largely unknown although an autoimmune origin is widely accepted. The response to immunomodulatory therapy, including intravenous immunoglobulins (IVIg), glucocorticosteroids and plasma exchange, supports the autoimmune hypothesis and the role of humoral factors, including autoantibodies, in its pathogenesis. Traditional CIDP pathogenic models describe the presence of combined cell-mediated and humoral immunity that result in an aberrant immune response targeting myelinated fibres of peripheral nerves^1^. However, the relative contribution of each component of the immune response is unknown. The recent discovery of disease-specific antibodies, such as anti-neurofascin 155 (anti-NF155), anti-contactin-1, anti-contactin-associated protein 1 and nodal neurofascin antibodies, that are only present in 5-10% of patients^2–4^, suggests the existence of small but homogeneous subgroups of CIDP patients in which specific effector mechanisms drive the disease. This model may explain better patient heterogeneity within the CIDP spectrum. Indeed the clinical, pathological and genetic heterogeneity disappears when patients are stratified according to highly specific biomarkers, such as autoantibodies^5–7^. The recent description of a significantly increased frequency of the HLA DRB1*15 allele in anti-NF155 antibody-positive patients in comparison with those anti-NF155 antibody-negative, an association that remained hidden before the description of these antibodies, supports this hypothesis^7^. These findings strongly suggest that, even though CIDP has an autoimmune pathogenesis, genetic factors could be essential in the development of CIDP in a subset of patients. Unfortunately, due to the rarity of the disease and the difficulty to recruit biologically homogeneous series of patients, research on the genetic factors related to CIDP is scarce^8^.

A non-synonymous homozygous mutation (p.Cys58Tyr) in CD59, a complement inhibitor present in the surface of red blood cells, was discovered in five Jewish children from North-Africa with chronic haemolysis and childhood relapsing immune-mediated polyneuropathy^9^. Interestingly, their symptoms were very similar to those of patients suffering from CIDP and the children had a partial response to IVIg and corticosteroid treatment.

We then hypothesized that *de novo* mutations in *CD59* might account for a fraction of adult-onset sporadic CIDP patients. With the aim of evaluating the possible role of *CD59* in CIDP, its coding region was fully sequenced in a homogeneous series of patients who did not present detectable anti-NF155, anti-contactin-1, NF140/186 or CASPR1 autoantibodies.

## Material and methods

### Patients, samples, protocol approvals and patient consents

Patients diagnosed with CIDP according to the European Federation of Neurological Societies/Peripheral Nerve Society (EFNS/PNS) diagnostic criteria^10^ provided written informed consent to participate and were included in the study according to a protocol approved by our Institution’s Ethics’ Committee. Whole blood was drawn in EDTA tubes and DNA extracted following standard protocols and stored until needed.

### Genetic studies

The entire coding region of *CD59* gene was amplified by polymerase chain reaction (PCR) and Sanger sequenced on an ABI 3100 automatic sequencer (Applied Biosystems, Foster City, CA, USA). Resulting electropherograms were visually analyzed using Sequencher software (Gene Codes Corp. Ann Arbor, MI, USA). Primer pair sequences and PCR conditions are available under request. In silico evaluation of potential deleterious effects of CD59 genetic variants was performed with the CADD-score (http://cadd.gs.washington.edu/), Mutation Taster (http://www.mutationtaster.org/), SIFT (http://sift.jcvi.org/), and Human Splicing Finder tool (http://www.umd.be/HSF3/index.html).

## Results

A total of 35 patients (57% male, mean age at inclusion 61 years old) were included in the study. Direct sequencing of all coding exons of *CD59* was performed. Only one variant was detected in one patient (Figure 1), a heterozygous guanine to alanine substitution (c.18G>A) which resulted in the synonymous change rs111771149 (p.Gly6Gly). According to the genome aggregation database (http://gnomad.broadinstitute.org/) this is a rare variant with an allele frequency of 0.003 in ExAC database and 0.001899 in gnomAD database in non-Finnish Europeans*. In silico* analysis of possible damaging consequences did not reveal any potential deleterious effect related to this genetic variant. The previously reported pathogenic p.Cys58Tyr mutation was not present in our patient cohort.

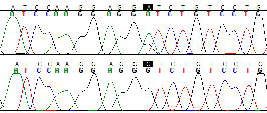

## Discussion

Our pilot study failed to identify functionally-relevant CD59 mutations in sporadic adult-onset CIDP patients suggesting that genetic dysfunction of CD59 is not a frequent cause of CIDP. A rare variant was found in one patient but was predicted to be functionally irrelevant.

*CD59* (protectin) encodes a glycosylphosphatidylinositol (GPI)-anchored cell surface membrane glycoprotein, which inhibits polymerization of complement molecule C9, the final step of membrane attack complex (MAC) formation, to protect host cells from complement-mediated lysis^9^.

Loss of function of *CD59* thus might makes cells, such as Schwann cells, susceptible to MAC-mediated lysis, possibly causing demyelination. In fact, complement deposits have been observed in areas of demyelination in sural nerve biopsies of patients with CIDP^11,12^. Also, *CD59*-deficient mice show MAC deposits in the perivascular tissue in the areas of demyelination, and are more susceptible to experimental autoimmune encephalomyelitis, inflammation and axonal loss than wild type mice ^13^.

Congenital *CD59* deficiency is an extremely rare and recently described disease characterized by hemolysis, recurrent ischemic strokes and relapsing immune-mediated polyneuropathy^9,14^.

After the description of the pathogenic *CD59* mutation p.Cys58Tyr^9^, other *CD59* mutations have been described. In 2015, an homozygous missense mutation (p.Asp49Val) was reported to be pathogenic in three Turkish patients from a two-generation family with immune-mediated peripheral neuropathy, chronic hemolysis and strokes^15^. Recently, a frameshift deletion in *CD59* (c.146delA, pAsp49Valfs*31) has been described in a 7-year-old girl with a demyelinating polyneuropathy and cerebral vasculopathy^14^.

These severe symptoms support the importance of *CD59* as an essential complement regulatory protein for protection of hematopoietic and Schwann cells against complement attack.

Acquired *CD59* deficiency is well known in paroxysmal nocturnal hemoglobinuria (PNH), where a clonal defect in GPI biosynthesis confined to the hematological system, results in hemolysis and prothrombotic tendency^16^. Eculizumab, a C5 inhibitor, is a treatment approved for PNH^17^. Therefore, treatment with eculizumab proved also effective in congenital CD59 deficiency; with a marked clinical improvement of these children, a reduction of hospitalizations and a reduction of IVIg and steroid doses^18^. Detection of CD59 mutations would have provided a novel therapeutic target in sporadic CIDP patients.

To our knowledge, this is the first genetic study aimed at identifying *CD59* alterations in CIDP. However, a study investigated the presence of antibodies targeting the complement inhibitors CD46, CD55 and CD59 and found no association of these antibodies with inflammatory neuropathies^19^.

In summary, our results suggest that CD59 mutations are not present in sporadic CIDP patients. We did not study somatic mutations restricted to nerve (mosaicism) or to regulatory regions of CD59 and, thus, the possibility that CD59 dysfunction plays a role in some CIDP patients has not been completely ruled out. Caution should be made in interpreting our data, since we used a limited series of patients. Since CIDP is a very rare disease and single-center case series are necessarily small, further genetic and immunologic studies in larger, collaborative, CIDP cohorts^20^ are needed to unravel the genetic basis of CIDP, improve pathogenetic models and find clinically useful biomarkers.

## Acknowledgments

The authors would like to acknowledge the Department of Medicine at the Universitat Autònoma de Barcelona. This project was supported by Fondo de Investigaciones Sanitarias (FIS), Instituto de Salud Carlos III, Spain and FEDER under grant FIS16/00627, and personal grant to LQ SLT006/17/00131 of the Pla Estratègic de Recerca i Innovació en Salut (PERIS) 2016-2020 of the Departament de Salut of the Generalitat de Catalunya, pricipal investigator Luis Querol.

